# Insecticide increases glyphosate resistance in *Ipomoea purpurea*

**DOI:** 10.64898/2025.12.20.695696

**Authors:** Grace M. Zhang, Regina S. Baucom

**Affiliations:** Ecology and Evolutionary Biology Department, University of Michigan

## Abstract

The evolution of plant resistance naturally occurs in complex, multifaceted environments that consist of multiple simultaneous stressors. Understanding how shifting environmental contexts may shape resistance evolution requires empirical studies that consider the combined effects of interacting potential stressors on fitness and selection. Here, we examined how exposure to an insecticide impacts the evolution of resistance to the herbicide glyphosate in *Ipomoea purpurea* (common morning glory). Through a factorial field experiment, we manipulated glyphosate and an insecticide to estimate selection on glyphosate and herbivory resistance. We found that glyphosate, not herbivory, acted as the primary agent of selection, favoring higher levels of glyphosate resistance. In the presence of glyphosate alone, positive correlational selection favored a combination of higher glyphosate and herbivory resistance, supporting prior work that suggested these traits may be linked. Importantly, insecticide exposure increased the level of glyphosate resistance and marginally weakened the strength of selection acting upon the trait. Together, our results indicate that the expression of herbicide resistance is context-dependent, and that additional chemicals like insecticide have the potential to alter the evolutionary trajectories of plant defense.

## INTRODUCTION

Plants experience natural and human-altered environments characterized by multiple, shifting stressors that may adversely impact plant physiology and/or fitness. How organisms adapt to such complex environments remains a central question in evolutionary biology. The evolution of plant resistance to damage—whether from natural enemies, abiotic stressors, or anthropogenic agents—provides some of the clearest demonstrations of adaptation in action. Classic studies of resistance have yielded insights into the genetic basis of adaptation, the costs of defense, and the trade-offs that constrain evolutionary trajectories (Warwick 1991; Bergelson and Purrington 1996; Strauss et al. 1999). For example, the framework of assessing fitness costs and benefits emerged from studies of plant defense against herbivory (Simms and Rausher 1987, 1989), and has since been extended to other naturally occurring biotic stressors such as fungal pathogens (Simms and Rausher 1993; Tian et al. 2003) and competition (Chaney and Baucom 2014), as well as abiotic stressors like herbicide (Baucom and Mauricio 2008; Debban et al. 2015).

Increasingly, studies recognize that organisms experience multiple, co-occurring stressors in nature (Siemens et al. 2003; Gassmann and Futuyma 2005; Wise and Rausher 2016), where both the benefits and costs of resistance can depend strongly on environmental context and shift across ecological gradients (Koricheva 2002; Cipollini et al. 2017). However, more empirical work involving multiple stressors is needed to predict plant evolutionary trajectories when selective pressures are indirect, nonlinear, or causally linked (Wise 2023).

Recent work has shown that overlapping biotic and abiotic stressors characterizing a complex environment may interact to shape both the expression and fitness consequences of plant defense (Côté et al. 2016; Desaint et al. 2021; Zandalinas et al. 2024). For example, in agricultural landscapes, anthropogenic chemicals such as herbicide and insecticide (potential abiotic stressors) are widely used to control pests (potential biotic stressors), which may impact plant evolutionary trajectories. Herbicides like glyphosate (the active ingredient of Roundup) are highly effective at weed control and impose strong selection on weed populations, driving the evolution of resistance (Culpepper 2006; Baucom 2019; Kreiner et al. 2022). Both the application of glyphosate and the evolution of glyphosate resistance may have cascading ecological and evolutionary consequences for plants and their associated communities, such as altering the patterns of insect herbivory (Zhang and Baucom 2024). At the same time, insecticides commonly co-occur with herbicides in agricultural contexts (Kwon and Penner 1995), potentially modifying the selective landscape beyond their direct effects on herbivores. Given that some insecticides such as cyantraniliprole are engineered to be taken up by plants to target piercing-sucking insects (Barry et al. 2015), and other insecticides like 2,4-D that are not meant to be systemic in plants can still be absorbed through foliar tissue (Finlayson and MacCarthy 1965), it is possible that insecticide itself may trigger plant defense pathways or response mechanisms, thus potentially interfering with the effects of herbicide. Indeed, combinations of herbicide and insecticide with diverse modes of action may differentially affect herbicide efficacy on plant species exposed to these chemicals (Daramola et al. 2023). However, while ecological interactions between herbicide and insecticide have been documented, there have been few empirical tests of whether the presence of a second chemical can alter the strength, direction, and causal pathways of selection on resistance traits.

We address this gap by examining how plant resistance evolves in environments mediated by anthropogenic chemicals: the herbicide glyphosate, which exerts strong selective pressure on plants, and the insecticide spinosad, which reduces insect herbivory. To test whether exposure to an insecticide influences the evolution of glyphosate resistance, we conducted a factorial common garden experiment examining the effects of each treatment environment on resistance and fitness in *Ipomoea purpurea* (common morning glory). Previous work in this species shows that glyphosate resistance confers a fitness benefit in the presence of glyphosate, but no detectable fitness cost measured as seed production in glyphosate’s absence (Baucom and Mauricio 2008; Van Etten et al. 2016). The addition of insecticide could modify these outcomes by interacting with glyphosate to strengthen or weaken the selective pressure it exerts on glyphosate resistance (Figure 1). For example, the insecticide spinosad has been shown to be systemic in *Solanum lycopersicum* (tomato) when applied to the roots, as it is able to control *Tetranychus urticae* (two-spotted spider mite) that are feeding on foliar tissue (Van Leeuwen et al. 2015). If *I. purpurea* is also able to uptake and translocate spinosad, then the presence of the insecticide in plant tissue may activate plant response mechanisms that could interfere with glyphosate’s effects. In this case, the fitness benefit of glyphosate resistance in the presence of glyphosate may be altered in strength by the application of the insecticide. Conversely, in the absence of glyphosate but in the presence of insecticide, glyphosate resistance may incur a cost, or alternatively confer a previously undetected fitness benefit.

**Figure 1.**
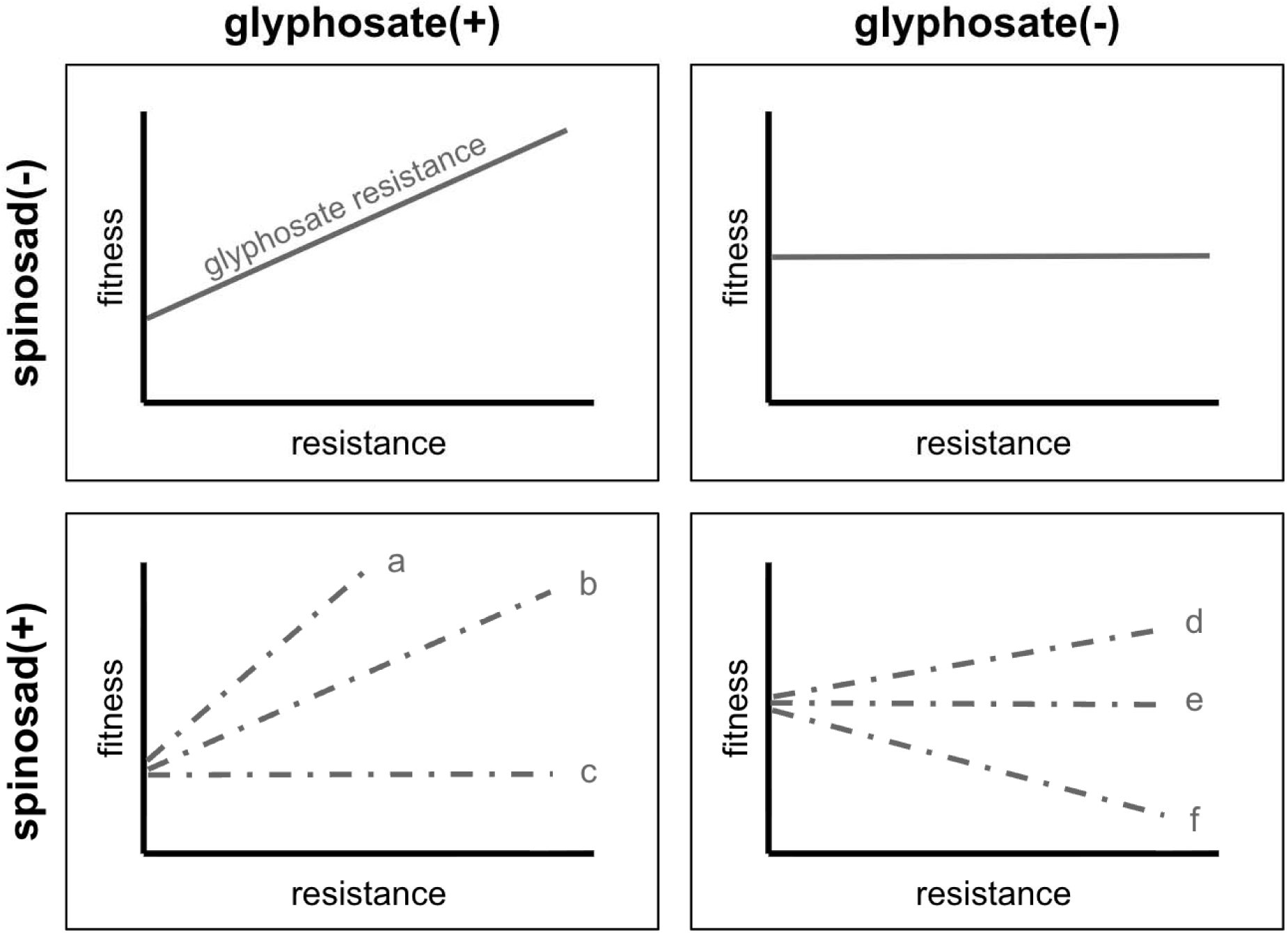
Factorial design of the common garden study, showing predictions of the fitness costs and benefits of glyphosate resistance in the form of selection analyses in the four treatment environments: glyphosate-present, spinosad-absent; glyphosate-absent, spinosad-absent; glyphosate-present, spinosad-present; and glyphosate-absent, spinosad-present. Solid lines represent relationships based on prior work in the system and dashed lines represent predictions made in this study. Based on past studies, there is a fitness benefit (glyphosate-present, spinosad-absent environment), but not a fitness cost (glyphosate-absent, spinosad-absent environment) in terms of seed production. We examine whether the addition of spinosad, an agriculturally relevant and commonly used insecticide, alters those selection patterns (as represented by the dashed lines in the two spinosad-present environments). The benefit of glyphosate resistance in the presence of glyphosate (glyphosate-present, spinosad-present environment) may be enhanced by spinosad such that it is greater than in the glyphosate-present, spinosad-absent environment (a), unaffected such that it is the same as in the spinosad-absent environment (b), or dampened such that it is less than in the spinosad-absent environment (c). Likewise, the cost of glyphosate resistance in the absence of glyphosate (glyphosate-absent, spinosad-present environment) may be outweighed by the benefit (d), remain undetectable like in the glyphosate-absent, spinosad-absent environment (e), or manifest (f). Spinosad as a secondary stressor may potentially interact with glyphosate to shape the evolutionary trajectory of glyphosate resistance. ALT TEXT: Four panels arranged in a factorial design with “glyphosate (+)” and “glyphosate (-)” on top, and “spinosad (+)” and “spinosad (-)” going up the left-hand side. In each panel is a line graph depicting a predicted selection analysis, with glyphosate resistance on the x-axis and fitness on the y-axis.

In this study, we explore how interacting anthropogenic chemicals—herbicide and insecticide— may influence plant adaptation by directly impacting fitness and indirectly *via* trait-mediated pathways by asking: 1) Is there a fitness benefit of glyphosate resistance (Figure 1 glyphosate+ spinosad–), but no evidence of a cost in terms of seed production (Figure 1 glyphosate– spinosad–), in line with previous work? 2) Are glyphosate and herbivory resistance under correlational selection, building upon prior research (Zhang and Baucom 2024)? If so, this suggests selection may favor a combination of the two forms of resistance that may not be evident simply from the detected costs and benefits (Svensson et al. 2021). 3) Does insecticide exposure impact glyphosate resistance directly and/or alter the patterns of selection acting on this trait? If so, the direction and/or magnitude of selection acting on glyphosate resistance in the presence of insecticide (Figure 1 spinosad+) would differ from the patterns of selection detected in spinosad’s absence (Figure 1 spinosad–). And 4) do glyphosate and insecticide application directly and/or indirectly impact the plant evolutionary trajectory *via* resistance to glyphosate or herbivory? Untangling direct selection from indirect selection is a key component of studying multivariate evolution, and a factorial experimental design allows for the examination of chains of selective causality in complex environments.

## MATERIALS AND METHODS

### Study system

*Ipomoea purpurea* (L.) Roth (Convolvulaceae), the common morning glory, is an annual weed frequently found in agricultural fields and disturbed soil beds in the United States. The species begins germination in the early spring and continues through the growing season. It grows rapidly, capable of producing over 8,000 seeds in a season before being killed by the first winter frost (Chaney and Baucom 2014). In Michigan, *I. purpurea* is fed upon by generalist coleopteran and lepidopteran herbivores such as *Popillia japonica* (Japanese beetle) and *Trichoplusia ni* (cabbage looper) (Zhang and Baucom 2024), and prior work shows that herbivory resistance is under positive selection in this species (Rausher and Simms 1989). Populations of the species are also naturally resistant to the herbicide glyphosate, likely through detoxification and cell transport mechanisms (Baucom and Mauricio 2004; Kuester et al. 2015; Gupta et al. 2023).

### Field experiment

On June 9, 2022, we planted replicate *Ipomoea purpurea* seeds from 45 family lines in a common garden field plot at the Matthaei Botanical Gardens in Ann Arbor, Michigan. These lines were generated from crosses between individuals (*i.e.*, maternal lines) sampled as seed from natural populations found in the US Southeast that exhibited variable levels of glyphosate resistance (Kuester et al. 2015). Individuals were randomly chosen from populations, selfed once, and then crossed in a haphazard design to randomize the genetic background. To generate experimental individuals, we selfed between one to five randomly selected half-sib families from 19 paternal lines to generate sufficient sample sizes. Four replicate seeds from each half-sib family were then randomly planted in each of four treatment environments in a factorial design of herbicide (glyphosate) and insecticide (spinosad, details below)—1) glyphosate-present, spinosad-absent, 2) glyphosate-absent, spinosad-absent, 3) glyphosate-present, spinosad-present, and 4) glyphosate-absent, spinosad-present. These combinations were repeated in two blocks, for a total of 1440 individual seeds. We watered daily and used a combination of weed fabric and hand-weeding to control weeds in all treatment environments. Germination started a few days after planting and was complete within ten days.

Two weeks after the seeds were planted, we started applying the field dose of 0.30 kg ai/L of the insecticide spinosad (Garden Insect Spray, Monterey Lawn and Garden Products, Inc., Fresno, CA, USA) to the plants using a handheld pump sprayer (Chapin International, Inc., Batavia, NY, USA) every other week for a total of five applications throughout the length of the experiment, as directed by the manufacturer. Spinosad, a broad-spectrum organic insecticide derived from a naturally occurring soil bacterium *Saccharopolyspora spinosa* (Bacteria: Actinobacteridae) (Mertz and Yao), is commonly used to protect vegetable crops and stored grain against insect herbivores by triggering uncontrollable muscle activity in targeted insects that leads to death (Mertz and Yao 1990; Bunch et al. 2014; Zhang et al. 2021). We conducted a preliminary growth room experiment to ensure that spinosad would not affect plant biomass on its own or by interacting with glyphosate (see Figure S1 for details). We counted the number of leaves on all plants five weeks after planting in order to estimate plant size (Debban et al. 2015). Six weeks after planting and before the plants started flowering, we applied the herbicide glyphosate (Roundup WeatherMax, Bayer Corp., Whippany, NJ, USA) at the sublethal dose of 1.0 kg ai/ha to the glyphosate-present treatment plots using a handheld CO_2_-pressurized sprayer (Spraying Systems Co., Wheaton, IL, USA) at a rate of 187 L/ha at 30 psi with a stride pace of 90 paces/min (Kuester et al. 2015). Because it was necessary to measure individual-level fitness later in the experiment, we chose a dose lower than the recommended field dose for *Ipomoea spp.* of 1.54 kg ai/ha (Monsanto 2017). Our chosen dose of 1.0 kg ai/ha was in line with past studies that aimed to merely stress the plants rather than eradicate them and is well within the range of recommended field doses of glyphosate for weed control (Duke et al. 2012; Vázquez et al. 2021). We scored damage two weeks after glyphosate application by dividing the number of damaged (*i.e.,* yellowed, browned, and/or shriveled) leaves on each plant by the total number of leaves in order to calculate the proportion of damaged leaves (per (Baucom and Mauricio 2008)). We also took note of which plants had died due to glyphosate application.

Five weeks after the glyphosate application, we randomly collected four leaves from each plant for herbivory damage estimates. Leaves were sandwiched between paper towels and kept cool on ice to be brought back to the lab, where they were scanned with a Canon CanoScan LIDE 400 flatbed scanner (Canon USA, Melville, NY, USA) to generate high-resolution images at 600 dpi. Herbivory damage was estimated as the percentage of tissue missing per leaf using the iOS app LeafByte (Getman-Pickering et al. 2020), then averaged across the four leaves for each plant. In line with Zhang and Baucom (2024), we elected to be conservative with our herbivory damage estimates and excluded ambiguous damage that could be attributed to glyphosate from our estimates. Additionally, we noticed that many plants had leaves exhibiting patches of rust fungus; thus, we scored the presence/absence of the fungus on each leaf as a binomial (1 -present, 0 -absent) and averaged those values across the four leaves for each plant. Finally at the end of the field season, we collected all seeds that were present on each plant in two thorough rounds in order to estimate fitness using seed number.

### Statistical analyses

We operationally defined resistance as 1-*p*, where *p* is the amount of damage experienced by an individual plant. Thus, when calculating glyphosate resistance, *p* refers to the proportion of leaves damaged by glyphosate per plant (Baucom and Mauricio 2008). Likewise, when calculating herbivory resistance, *p* is the average percentage of leaf tissue missing due to herbivory for each plant (Rausher and Simms 1989).

All analyses were conducted using R 4.3.2 (R Core Team 2023). For the field experiment, we excluded any individual plants that did not germinate (N = 229), as well as plants in the glyphosate-absent treatments that were inadvertently damaged by glyphosate (N = 43). We also removed any plants that died before herbivory data could be recorded. All surviving plants were included in the analyses, including plants that produced no seeds. Ultimately, we had 292 plants in the glyphosate-present, spinosad-absent treatment, 264 plants in the glyphosate-absent, spinosad-absent treatment, 294 plants in the glyphosate-present, spinosad-present treatment, and 279 plants in the glyphosate-absent, spinosad-present treatment, for a total of 1129 plants.

### Treatment effects of glyphosate and spinosad

We wanted to confirm the efficacy of treatment regimes. First, we built linear mixed models using the package *lme4* (Bates et al. 2015) to determine if spinosad reduced herbivory as expected. Using all experimental plants, we constructed the following model:

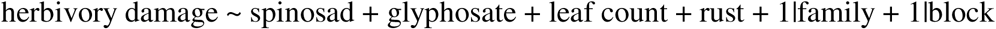

We transformed the response variable of herbivory damage using rank normalization in order to ensure normality after using the package *bestNormalize* (Peterson 2021) to identify the appropriate transformation. Spinosad and glyphosate were included as fixed effects, while leaf count (as a proxy for plant size) and the proportion of collected leaves showing rust fungus were included as covariates in all models. The Satterthwaite approximation was used to determine the degrees of freedom of the fixed-effect variables. Family line and block were included as random effects unless stated otherwise. To test the significance of random effects, we removed each one and used the χ^2^ test of difference (df = 1) to compare the simplified model to the full model.

Next, we wanted to determine if glyphosate influences herbivory damage (based on prior results; Zhang and Baucom 2024) or alternatively, if spinosad altered glyphosate resistance. We first tested for an effect of glyphosate application on herbivory damage in the spinosad-present and -absent environments separately using the following model, with herbivory damage transformed using rank normalization in both treatment environments:

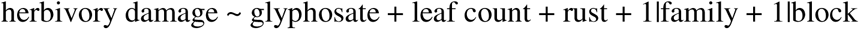

To examine whether spinosad application may have had an effect on glyphosate resistance, we built the following linear mixed model using just the plants treated with glyphosate, with glyphosate resistance transformed using rank normalization:

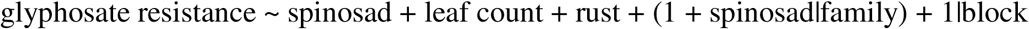

We specifically tested for an interaction between spinosad and family line while letting the slopes and intercepts vary because we were interested in whether the effect of spinosad application on glyphosate resistance showed genetic variation.

### Genetic variation and among-family variance in glyphosate and herbivory resistance

We assessed if there was evidence for genetic variation for glyphosate and herbivory resistance by including family line as a fixed effect in the following linear mixed model:

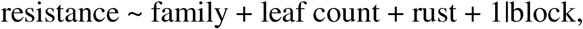

where the resistance was either glyphosate or herbivory resistance. For resistance to either potential stressor (*i.e.*, glyphosate and herbivory), we tested for genetic variation first among all plants that had been exposed to that stressor to determine overall variation in the trait, then in both the presence and the absence of the other stressor to account for any treatment effects of the second stressor. Glyphosate resistance as a response variable in the presence of spinosad was transformed using the Yeo-Johnson transformation to ensure normality, while both glyphosate and herbivory resistance in all other treatment environments were transformed using rank normalization.

Using the model we previously built to assess the effect of spinosad on glyphosate resistance (glyphosate resistance ∼ spinosad + leaf count + rust + (1 + spinosad|family) + 1|block), we also determined whether the presence of spinosad affected the partitioning of variance in glyphosate resistance attributable to family line among glyphosate-treated plants. This model allowed for among-family variance to be partitioned separately depending on the presence of spinosad, with variance in the intercept reflecting among-family variance in the spinosad-absent environment, and variance in the slope reflecting additional among-family variance attributed to differential family responses to spinosad. We then used parametric bootstrapping (n = 1000 simulations) to generate 95% confidence intervals on intraclass correlation coefficients (ICCs) within spinosad environments. ICCs were calculated in each treatment environment as the ratio of among-family variance to total variance (*i.e.*, among-family + block + residual variance).

Similarly, we tested if the presence of glyphosate influenced the variance partitioning in herbivory resistance attributable to family line among plants in the spinosad-absent environment using a similar model: herbivory resistance ∼ glyphosate + leaf count + rust + (1 + glyphosate|family) + 1|block. ICCs and 95% confidence intervals were calculated in the presence and absence of glyphosate as previously described.

### Selection analyses on resistance traits

We conducted selection analyses on the two forms of resistance in each of the four treatment environments to determine if either spinosad or glyphosate would affect the patterns of selection acting on resistance to glyphosate and herbivory. We also assessed if there were fitness costs and benefits of each form of resistance depending on the treatment environment. If there was positive selection for either glyphosate or herbivory resistance in the presence of the corresponding potential stressor, then we would conclude there was a fitness benefit associated with that form of resistance. The converse was also examined—negative selection for a form of resistance in the absence of the stressor would suggest there was a fitness cost associated with resistance to that stressor.

In accordance with past studies, we used a modified form of Lande and Arnold’s method of detecting selection using partial regression coefficients (Lande and Arnold 1983; Rausher 1992; Mauricio and Mojonnier 1997). Instead of performing selection analyses on phenotypic trait values, we used family line means of trait values to conduct genotypic selection analyses in order to eliminate environmental biases (Rausher 1992; Stinchcombe et al. 2002). Additionally, we used the residuals of untransformed values of relative fitness and resistance after removing the effects of block to further remove environmental effects from the analyses. Relative fitness was calculated within each treatment environment by dividing the fitness of each plant by the average fitness of all plants in that environment. After calculating family means of relative fitness and resistance trait values, we standardized the resistance traits in each treatment environment such that they had an average of zero and a standard deviation of one. Relative fitness was then regressed on each form of resistance within each treatment environment to determine the pattern of selection for that particular trait. Directional selection was estimated using models with only linear terms, while nonlinear (*i.e.*, stabilizing/disruptive) selection was estimated using models containing both linear and quadratic terms. All selection gradients for nonlinear selection were reported as twice the quadratic regression coefficient.

In the glyphosate-present, spinosad-absent environment (Figure 1), we assessed the fitness benefits of glyphosate and herbivory resistance. The parentheses next to each trait indicates the treatment environment from which mean trait values were obtained, with plus and minus signs signifying the presence and absence of either chemical in line with Figure 1:

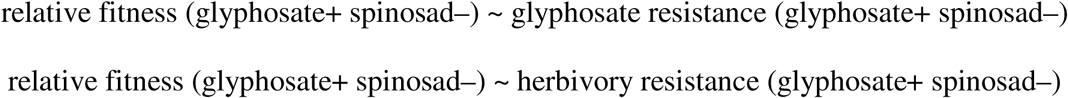

In the glyphosate-absent, spinosad-absent environment, we tested for the cost of glyphosate resistance and the benefit of herbivory resistance:

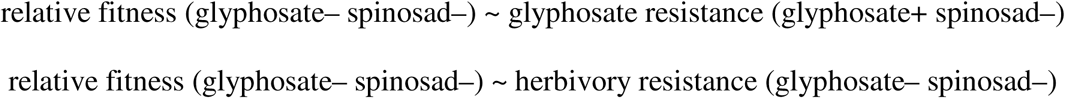

In the glyphosate-present, spinosad-present environment, we assessed the benefit of glyphosate resistance and cost of herbivory resistance:

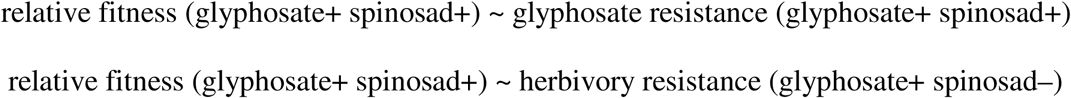

Lastly, in the glyphosate-absent, spinosad-present environment, we tested for the costs of glyphosate and herbivory resistance:

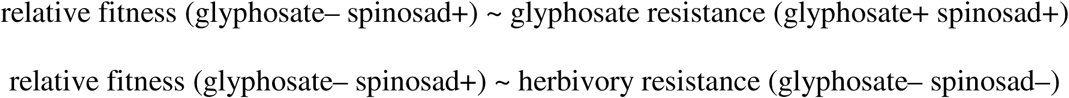

To detect correlational selection between the two forms of resistance in each treatment environment, we regressed relative fitness on the interaction term between the two resistance traits, as well as all linear and quadratic terms of the resistance traits. We then used the *fields* package (Nychka et al. 2021) to generate a fitness landscape fitted with thin-plate splines. We further tested for genetic correlations between the two forms of resistance in each treatment environment by calculating family-level Pearson’s correlation coefficients.

Since we were interested in whether spinosad might alter patterns of selection (if any) that were acting on glyphosate resistance, we built the following model to compare the selection gradients between the two glyphosate-present environments:

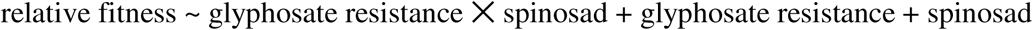

Relative fitness and the standardization of glyphosate resistance were calculated within treatment environments, as described above. A significant interaction between glyphosate resistance and spinosad would suggest that the selection gradients were altered by the presence of spinosad.

### Direct and indirect effects on fitness

We assessed the direct and indirect effects of glyphosate and spinosad on fitness—how do the dual treatments of herbicide and insecticide affect resistance, and how does that impact fitness? To answer these questions, we constructed an SEM using all experimental plants, specifically testing the effects that glyphosate and spinosad have on both glyphosate resistance and herbivory resistance, as well as seed number. We also tested for relationships between the resistances and fitness. With family line and block included as random effects, the following relationships were modeled using the package *piecewiseSEM* (Lefcheck 2016), and the Fisher’s C statistic was used to determine model fit:

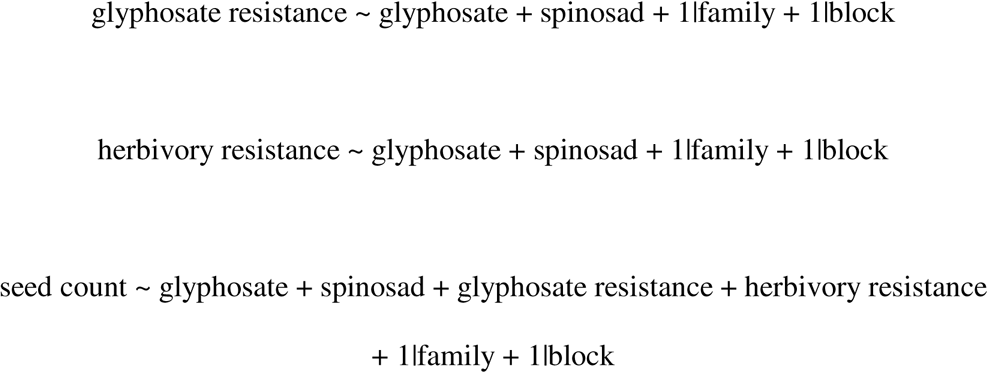

We did not include a relationship between glyphosate resistance and herbivory resistance in the final model. Doing so would have resulted in a saturated model, which would not have allowed us to assess model fit. Additionally, the saturated model reported that the relationship between the two resistances was not significant, and including the relationship resulted in a model with a higher AIC score (saturated: 7179.76 vs. current model: 7173.24), which suggests a worse fit.

Thus, we proceeded with the SEM that omitted this relationship.

## RESULTS

### Treatment effects of glyphosate and spinosad

Glyphosate was an effective stressor of *Ipomoea purpurea*. While only ten plants died from herbicide application, all but one of the 586 glyphosate-treated plants exhibited leaf damage from glyphosate (on average, 37.3% of leaves per plant were damaged) and reduced total seed production (mean seed number in glyphosate-treated plants: 286 vs. untreated plants: 1714).

Overall, plants exposed to glyphosate experienced more chewing herbivory damage than unexposed plants (glyphosate-present: 0.37% per plant, glyphosate-absent: 0.29% per plant; F□,□□□□ = 4.90, p = 0.027; Figure 2a, Table S1). This effect held when spinosad was present (glyphosate-present: 0.18% per plant, glyphosate-absent: 0.12% per plant; F _,542_ = 5.32, p = 0.021; Table S2), but not when spinosad was absent (F□,□□□□ = 1.18, p = 0.278; Table S2).

**Figure 2.**
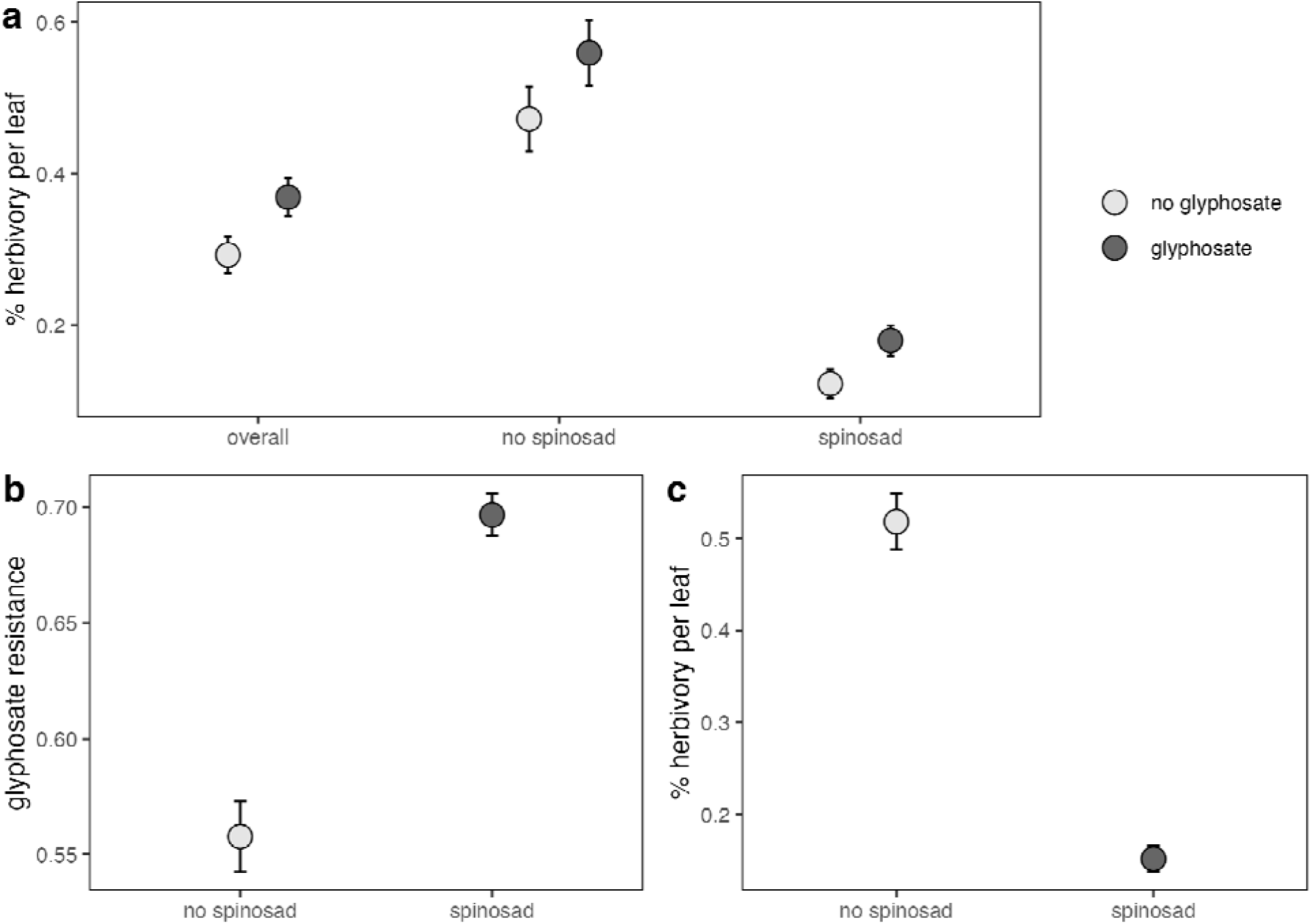
Treatment effects of glyphosate and spinosad application. Circles show unadjusted treatment means with standard error bars. (a) Plants treated with glyphosate experienced a higher level of herbivory damage overall (F_1,1103_ = 4.90, p = 0.027), as well as in the presence of spinosad (F_1,542_ = 5.32, p = 0.021). In the absence of spinosad, however, glyphosate did not affect herbivory damage (F_1,551_ = 1.18, p = 0.278). (b) Spinosad-treated plants had higher levels of glyphosate resistance (F_1,227_ = 82.17, p < 0.001). (c) Spinosad-treated plants experienced less herbivory damage (F_1,1095_ = 198.94, p < 0.001). ALT TEXT: Three-panels of plots showing the treatment effects of glyphosate and Spinosad on herbivory damage and glyphosate resistance.

Plants that were exposed to both glyphosate and spinosad showed higher levels of resistance to glyphosate than plants that were just exposed to glyphosate (spinosad: 69.67% leaves showed no glyphosate damage, no spinosad: 55.77% leaves showed no glyphosate damage; F_1,227_ = 82.17, p < 0.001; Figure 2b, Table S3). There was no interaction between spinosad treatment and family line, however, indicating that families generally experienced similar levels of increased glyphosate resistance in the presence of the spinosad (χ^2^ = 1.23, p = 0.540; Table S3). Spinosad application significantly reduced herbivory overall (untreated: 0.52%, treated: 0.15%; F_1,1095_ = 198.94, p < 0.001; Figure 2c, Table S1), but did not eliminate herbivory.

### Genetic variation and among-family variance in resistance to glyphosate and herbivory

We investigated whether there was genetic variation for the two forms of resistance and found evidence of genetic variation for glyphosate resistance across all plants in glyphosate-present environments (F_44,538_ = 1.72, p = 0.003; Table S4). This was also the case in both the absence of spinosad (F_44,244_ = 1.77, p = 0.004; Table S4) and in the presence of spinosad (F_44,246_ = 1.73, p = 0.005; Table S4). Among-family variance in glyphosate resistance in the presence of spinosad (ICC = 0.101, 95% CI: 0.028 -0.200; Table S5) was comparable to in spinosad’s absence (ICC = 0.049, 95% CI: 0.007 -0.130; Table S5) given the wide, overlapping confidence intervals.

In contrast to glyphosate resistance, there was no evidence of genetic variation underlying herbivory resistance among plants that were not treated with spinosad (F_44,508_ = 0.59, p = 0.984; Table S6). Nor was there evidence of genetic variation for herbivory resistance in either the glyphosate-present or glyphosate-absent environments (glyphosate-present: F_44,245_ = 0.57, p = 0.987; glyphosate-absent: F_44,216_ = 1.04, p = 0.414; Table S6). Likewise, there was little difference in among-family variance in herbivory resistance in the presence and absence of glyphosate, given the largely overlapping confidence intervals (glyphosate-present ICC = 0.001, 95% CI: 0.000 -0.078; glyphosate-absent ICC = 0.004, 95% CI: 0.000 -0.082; Table S7).

### Glyphosate resistance is under positive selection

We tested for selection acting upon family line averages of glyphosate and herbivory resistance in each of the four treatment environments in order to determine if the presence or absence of either glyphosate or spinosad would alter the patterns of selection, as well as whether there were fitness costs and benefits associated with either form of resistance. Significant selection was detected in two environments. In the presence of glyphosate and the absence of spinosad (*i.e.*, herbivores were present), there was positive selection for glyphosate resistance, indicating a fitness benefit of this trait (β = 0.46, F_1,43_ = 28.56, p < 0.001; r = 0.61, p < 0.001; Figure 3a; Table 1). Similarly, in the presence of both glyphosate and spinosad, there was positive selection for glyphosate resistance (β = 0.24, F_1,43_ = 6.15, p = 0.017; r = 0.35, p = 0.017; Figure 3c; Table 1). The interaction between spinosad and glyphosate resistance on fitness was marginally significant (F_1,86_ = 2.92, p = 0.091; Table S8), suggesting that spinosad may have weakened selection acting on glyphosate resistance (β = 0.46 without spinosad vs. 0.24 with spinosad)..

**Figure 3.**
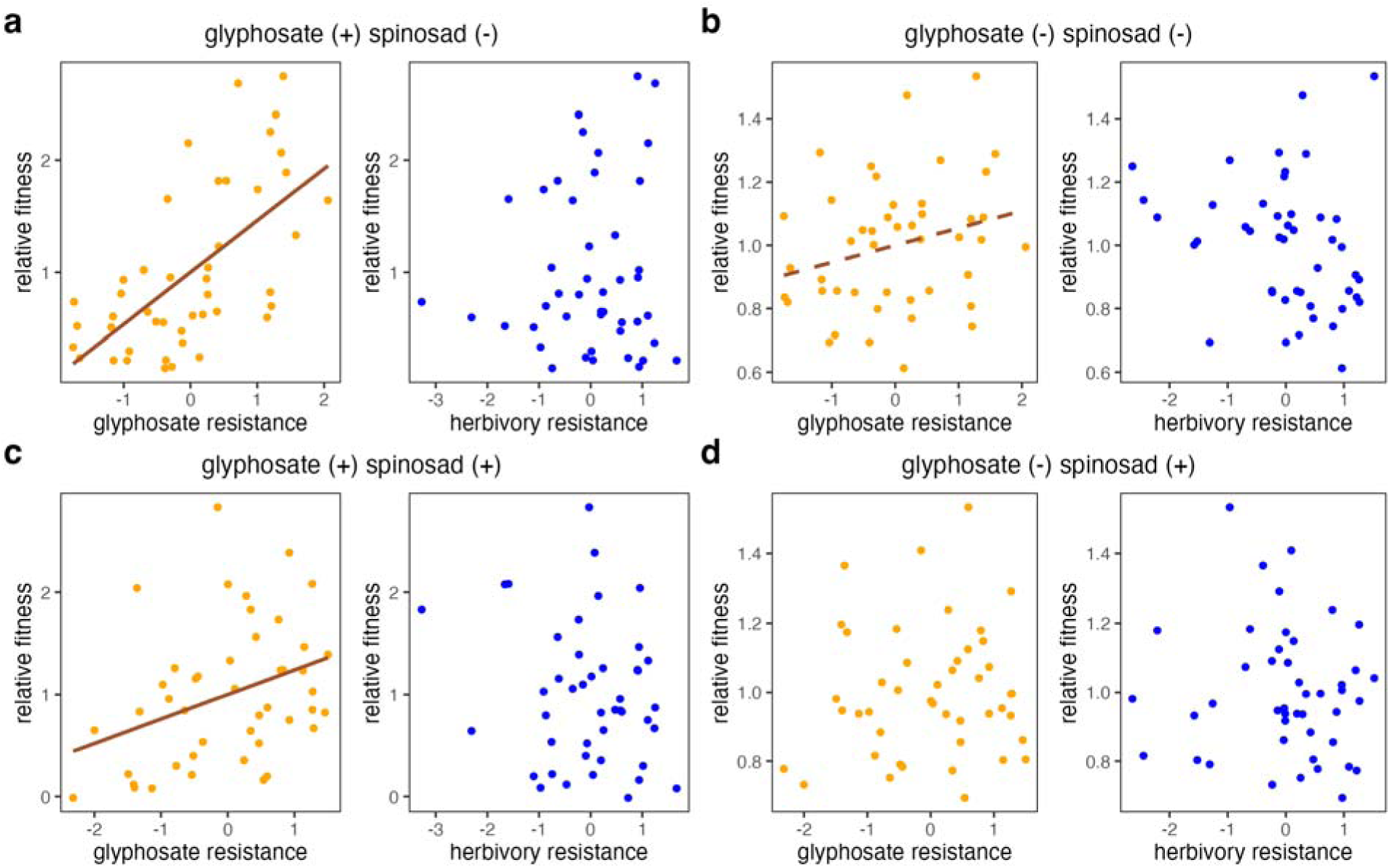
Family means of fitness in relation to resistance to glyphosate (orange) and herbivory (blue) in the four treatment environments. Each point is a family-line mean for that particular trait. (a) In the glyphosate-present, spinosad-absent environment, there was positive selection for glyphosate resistance (β = 0.46, F_1,43_ = 28.57, p < 0.001), as denoted by the solid line. (b) In the glyphosate-absent, spinosad-absent environment, there was marginal positive selection acting on glyphosate resistance (β = 0.05, F_1,43_ = 3.24, p = 0.079), as denoted by the dashed line. (c) In the glyphosate-present, spinosad-present environment, there was positive selection for glyphosate resistance (β = 0.24, F_1,43_ = 6.15, p = 0.017) as denoted by the solid line. (d) In the glyphosate-absent, spinosad-present environment, there was no selection acting on either form of resistance. ALT TEXT: Four panels represent the treatment environments in the field study. In each panel, there are two scatterplots, with one depicting a selection analysis for glyphosate resistance and the other showing a selection analysis for herbivory resistance.

**Table 1.** Selection analyses for resistance to glyphosate and herbivory in each of the four treatment environments. Models for each type of resistance were conducted separately in each environment by regressing relative fitness on the standardized form of each resistance. Bolded p-values denote statistical significance, while italicized p-values show marginal statistical significance. relative fitness ∼ standardized resistance

| Trait | $\beta$ | SE | F | df | p |
| --- | --- | --- | --- | --- | --- |
| *glyphosate-present, spinosad-absent* |  |  |  |  |  |
| glyphosate resistance | 0.46 | 0.09 | 28.57 | 1 | **<0.001** |
| herbivory resistance | 0.05 | 0.11 | 0.21 | 1 | 0.652 |
| *glyphosate-absent, spinosad-absent* |  |  |  |  |  |
| glyphosate resistance | 0.05 | 0.03 | 3.24 | 1 | *0.079* |
| herbivory resistance | -0.05 | 0.03 | 2.52 | 1 | 0.119 |
| *glyphosate-present, spinosad-present* |  |  |  |  |  |
| glyphosate resistance | 0.24 | 0.10 | 6.15 | 1 | **0.017** |
| herbivory resistance | -0.12 | 0.10 | 1.38 | 1 | 0.246 |
| *glyphosate-absent, spinosad-present* |  |  |  |  |  |
| glyphosate resistance | 0.01 | 0.03 | 0.26 | 1 | 0.616 |
| herbivory resistance | -0.02 | 0.03 | 0.30 | 1 | 0.587 |

There was also marginally significant positive selection for glyphosate resistance detected in the absence of both glyphosate and spinosad (β = 0.46, F_1,43_ = 3.24, p = 0.079; r = 0.26, p = 0.079; Figure 3b; Table 1). No other linear or quadratic selection on glyphosate or herbivory resistance was detected in other treatment environments (Figures 3d; Tables 1 and S9).

Further, in the treatment environment with only glyphosate, there was correlational selection acting on resistance to glyphosate and herbivory, favoring high levels of both forms of resistance (correlational selection gradient = 0.21, F_1,39_ = 6.45, p = 0.015; Figure 4; Table S9). However, there was no selection acting on herbivory resistance in this environment (β = 0.05, F_1,43_ = 0.21, p = 0.652; Figure 3a; Table 1), and the two resistance traits were not genetically correlated (r = 0.04, p = 0.794). There were no signs of correlational selection in any other treatment environment (Table S9).

**Figure 4.**
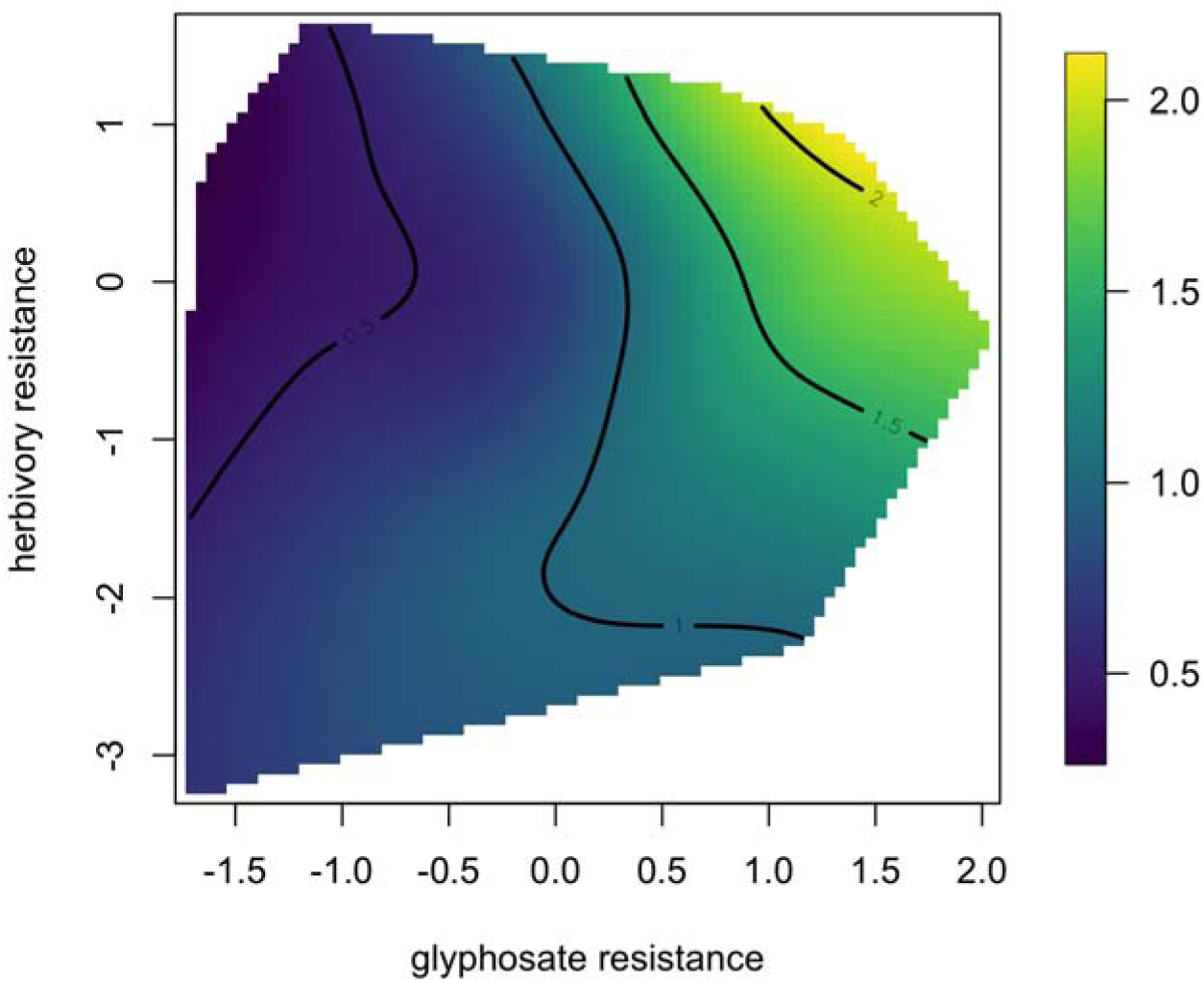
In the glyphosate-present, spinosad-absent environment (*i.e.*, herbivores are present), there is correlational selection that favors high levels of both glyphosate and herbivory resistance (correlational selection gradient = 0.21, F_1,39_ = 6.45, p = 0.015). The colored bar on the side shows relative fitness, with higher values being warmer and lower values being cooler. ALT TEXT: Contour plot with glyphosate resistance on the x-axis and herbivory resistance on the y-axis, showing a peak near the top right-hand corner of the plot.

### Direct and indirect effects of treatments on resistance and seed production

We assessed the direct and indirect effects of glyphosate and spinosad treatments on glyphosate resistance, herbivory resistance, and fitness by constructing a well-fit structural equation model (SEM) that had a Fisher’s C statistic of 0.006 (p = 0.997; Figure 5). Glyphosate treatment negatively impacted glyphosate resistance (standardized β = -0.75, df = 1096, p < 0.001), which is necessarily the case because unexposed plants would not have experienced glyphosate damage and therefore would have a resistance level of 1 in the SEM. In turn, glyphosate resistance was positively associated with seed production, meaning that more resistant plants produced more seeds (standardized β = 0.28, df = 647, p < 0.001). In contrast, the impact of glyphosate on seed production was negative (total effect = -0.76), but split into the primary direct effect (standardized β = -0.55, df = 960, p < 0.001) and an indirect effect that was mediated by glyphosate resistance (-0.75 * 0.28 = -0.21). Glyphosate had no effect on herbivory resistance (standardized β = -0.04, df = 1098, p = 0.122).

**Figure 5.**
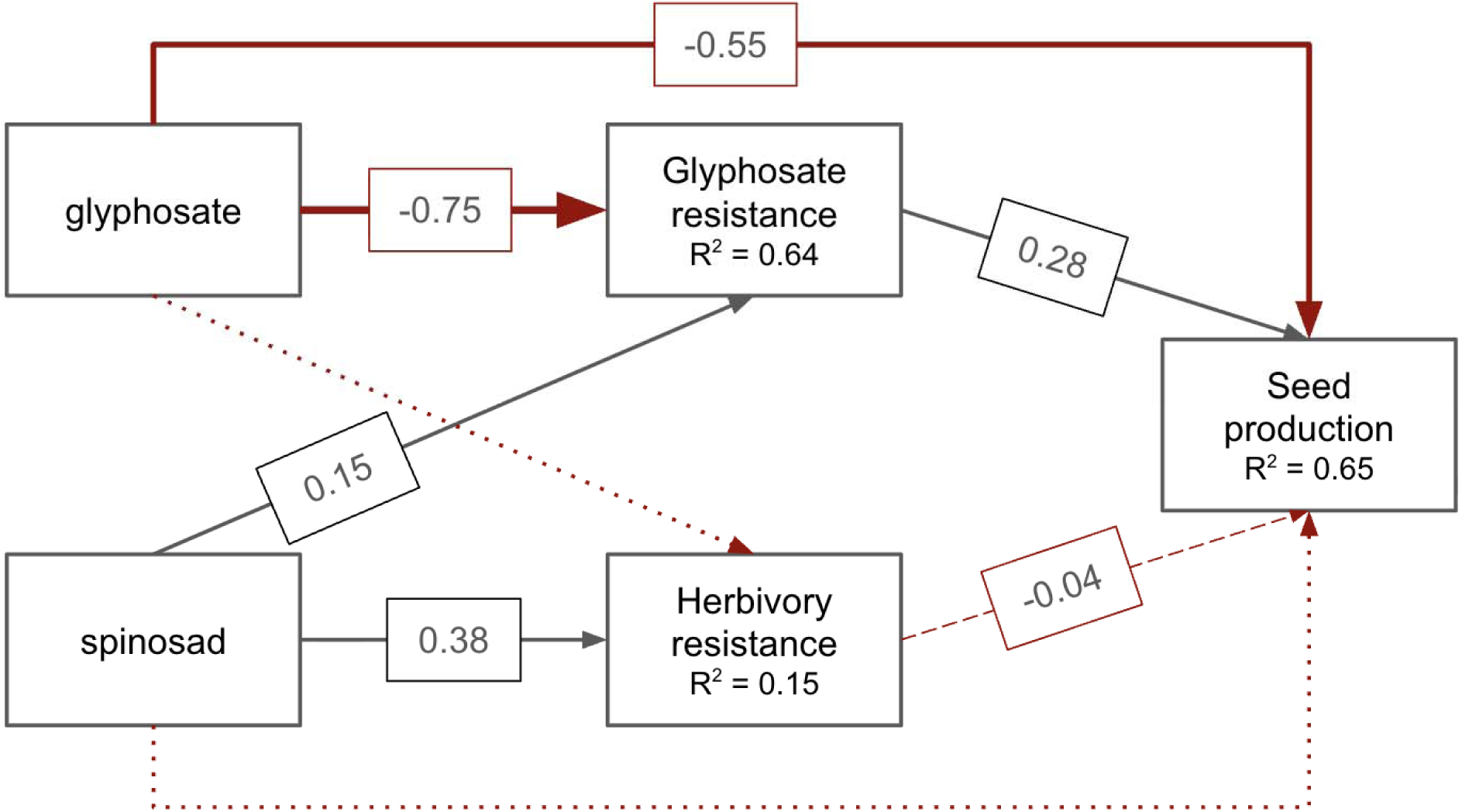
Structural equation model (SEM) of the relationships between glyphosate treatment, spinosad treatment, glyphosate resistance, herbivory resistance, and seed production. All models in the SEM are linear mixed models, with family line and block included as random effects. Boxes represent the measured variables and arrows show the unidirectional relationships between variables. Gray arrows indicate positive relationships and red arrows represent negative relationships. The thickness of the arrows reflect the magnitude of the standardized β coefficients, which are shown along the arrows. All solid arrows show statistically significant paths (p < 0.001), the dashed line denotes a marginally significant path, and dotted arrows show nonsignificant paths. Conditional R^2^ values are included within the box of each response variable. This model fit well, with a Fisher’s C score of 0.006 (p = 0.997). ALT TEXT: Diagram with glyphosate, glyphosate resistance, spinosad, herbivory resistance, and seed production in boxes. Solid red arrows point from glyphosate to glyphosate resistance and seed production, and a dotted red arrow points from glyphosate to herbivory resistance. Solid gray arrows point from spinosad to glyphosate resistance and herbivory resistance, and a dotted red arrow points from spinosad to seed production. A solid gray arrow points from glyphosate resistance to seed production. A red dashed arrow points from herbivory resistance to seed production.

Spinosad treatment led to increased herbivory resistance (*i.e.*, decreased herbivory; standardized β = 0.38, df = 1098, p < 0.001)—plants exposed to spinosad are by necessity more resistant to herbivory because they experience less herbivory damage. Spinosad also increased glyphosate resistance (standardized β = 0.15, df = 1096, p < 0.001), but had no direct effect on seed production (standardized β = -0.01, df = 1077, p = 0.708). Herbivory resistance had a marginally significant negative impact on seed production (standardized β = -0.04, df = 1104, p = 0.057).

Additionally, the negative effect of glyphosate and the positive effect of spinosad on fitness were both mediated through resistance to glyphosate to some degree.

## DISCUSSION

We conducted a factorial field experiment to examine how exposure to an insecticide (spinosad) shapes the evolution of resistance to an herbicide (glyphosate) in *Ipomoea purpurea*. Two main patterns emerge. First, glyphosate is predictably the dominant selective agent, consistently reducing fitness and favoring higher resistance in glyphosate-present environments. Second, spinosad unexpectedly increases glyphosate resistance, which in turn indirectly increases fitness. Together, these results show that glyphosate resistance is plasticand that a change in the environmental context can alter the expression of resistance traits and—through those traits— plant fitness..

### Glyphosate resistance is under positive selection

Glyphosate application, even at a sublethal dose, imposed strong fitness consequences as expected, delaying development and substantially reducing seed production. Consistent with previous work in this system (Baucom and Mauricio 2008), glyphosate resistance in the presence of glyphosate experienced significant positive selection in the spinosad-absent environment, indicating a clear fitness benefit of increased glyphosate resistance. Importantly, we detected genetic variation for glyphosate resistance across plants exposed to glyphosate, suggesting continued evolutionary potential (Baucom and Mauricio 2008). The structural equation model (SEM) reinforces this conclusion—glyphosate reduced fitness primarily through a strong direct effect, with an additional indirect effect mediated by glyphosate resistance. These results highlight glyphosate as the principal driver of the evolutionary trajectory of glyphosate resistance in this study. However, despite strong selection, we detected no fitness costs of glyphosate resistance measured as seed production, regardless of the presence of spinosad. This absence of detectable costs is consistent with prior studies that identified costs in seed viability, germination, or plant size (Debban et al. 2015; Van Etten et al. 2016) but not seed production.

Although glyphosate resistance clearly enhances fitness in the presence of herbicide, its associated costs may be conditional on dose, genetic background, and experimental environment (Neve et al. 2014). Further, our experimental design is only suitable to explicitly detect constitutive resistance that is always present, but not induced resistance triggered by glyphosate exposure (Van Zandt 2007). However, previous work has shown that glyphosate resistance in *Ipomoea purpurea* is likely to be constitutive (Gupta et al. 2023); although if the mechanism of resistance included induced resistance, we would not expect to identify fitness costs of resistance in the absence of glyphosate.

Interestingly, although there was no univariate selection on herbivory resistance, we did find correlational selection favoring jointly higher levels of glyphosate and herbivory resistance in the presence of glyphosate and absence of spinosad (*i.e.*, herbivores were present). This pattern suggests that as glyphosate resistance evolves, herbivory resistance may also increase—even in the absence of detectable genetic variation for herbivory resistance itself. This interpretation aligns with previous SEM-based work, which inferred a causal relationship between glyphosate and herbivory resistance (Zhang and Baucom 2024).

One explanation is that the two resistance traits may share underlying genetic mechanisms. Glyphosate inhibits 5-enolpyruvylshikimate-3-phosphate synthase (EPSPS), a key enzyme upstream of metabolic pathways that synthesize aromatic amino acids—the precursors of many secondary metabolites involved in chemical defense against herbivores (Amrhein et al. 1980; Mithöfer and Boland 2012; Fuchs et al. 2021). The likely mechanism of glyphosate resistance in *Ipomoea purpurea*—metabolic detoxification coupled with enhanced transport (Gupta et al. 2023)—could affect herbivory resistance in two ways. First, detoxification may reduce the number of glyphosate molecules that reach and inhibit EPSPS (Baek et al. 2021), thereby mitigating glyphosate’s indirect suppression of anti-herbivore defense compound biosynthesis. Second, loci underlying detoxification pathways (*e.g.*, cytochrome P450s, glycosyltransferases, and ABC and phosphate transporters) may have pleiotropic effects on herbivory resistance. For example, uridine diphosphate glucosyltransferases (UGTs) regulate the glycosylation of phytohormones central to herbivory defense such as abscisic acid, jasmonates, and salicylic acid (Dimunová et al. 2022; Gharabli et al. 2023). If glyphosate and herbivory resistance are linked due to pleiotropy, the relationship between the traits will persist, and correlational selection will likely shape the evolution of those traits concurrently over time (Svensson et al. 2021).

### Spinosad increases glyphosate resistance

One of the more interesting findings of this study was that spinosad application increased glyphosate resistance among glyphosate-exposed plants and indirectly increased fitness through this effect. . While our design does not allow us to fully disentangle the effects of spinosad from reduced herbivory, the SEM suggests that spinosad acts directly on glyphosate resistance rather than indirectly through herbivory. A plausible explanation for this effect is physiological priming *via* shared detoxification pathways. Resistance to spinosad in insects can be conferred *via* metabolic detoxification, particularly by upregulating enzymes such as monooxygenases, cytochrome P450s, and UGTs (Sparks et al. 2012; Bastarache et al. 2023; Wang et al. 2024).

Notably, plants have metabolic detoxification processes that trigger the same classes of enzymes (Edwards et al. 2011; Gharabli et al. 2023), and importantly, the same processes likely confer resistance to glyphosate in *Ipomoea purpurea* (Gupta et al. 2023). Thus, it is possible that the application of spinosad may prime plants to become more resistant to glyphosate by activating the same detoxification pathways. Moreover, while spinosad is generally thought to have low systemic activity in plants (National Organic Standards Board Technical Advisory Panel 2002), one study found that applying spinosad to the roots of *Solanum lycopersicum* (tomato) was enough to control the herbivore *Tetranychus urticae* (two-spotted spider mites) on its leaves (Van Leeuwen et al. 2005). Additionally, spinosad was detected in both the aboveground and belowground tissue of *Allium cepa* (onion) that had been treated with the insecticide as seeds for up to 60 days post planting (Moretti et al. 2021). It seems evident that spinosad has the potential to translocate in plant tissue. If *I. purpurea* can similarly take up spinosad, the insecticide may activate detoxification pathways, priming the plants to respond quickly to the herbicide and better protect from damage.

The potential of insecticides to modify herbicide efficacy has long been considered, given the potential for antagonizing chemicals to reduce weed control. However, such studies generally focus on herbicide performance in terms of plant death or reduction in biomass, without considering further ecological or evolutionary implications (Pankey et al. 2004; Daramola et al. 2023). Our work explicitly explores the consequences of insecticide application on the potential evolutionary trajectory of herbicide resistance in an agricultural weed, showing that spinosad enhances glyphosate resistance and thereby increases fitness. Additionally, we detected a marginally significant effect of spinosad on the positive selection acting on glyphosate resistance—indeed, the selection coefficient in the presence of spinosad is halved as compared to spinosad’s absence. This potential reduction of selective pressure may be caused by the increased glyphosate resistance in spinosad-exposed plants—the level of glyphosate resistance may be closer to the phenotypic optimum of the trait; thus, glyphosate would exert less selective pressure on the spinosad-exposed plants than not-exposed plants (Arnold 1992). Over time, plants exposed to both glyphosate and spinosad may reach the limit of glyphosate resistance that can evolve in this environment, although that limit may be lower than complete resistance.

Additionally, while these effects are likely species-specific, to our knowledge our study is the first to show that spinosad interacts with glyphosate to reduce herbicide efficacy in a weed (Sparks et al. 2004; Scroggs et al. 2005). Understanding how the interactions between herbicide and insecticide impacts the expression of herbicide resistance and fitness is essential for predicting the persistence of resistance in heterogeneous, complex environments such as agricultural landscapes.

### Context-dependent ecological interactions

Much of the work examining interactions between insecticides and herbicides have focused on the impact of insecticide on herbicide efficacy. However, herbicide may influence insecticide efficacy as well. This relationship has primarily been studied in lab settings, with an emphasis on insect mortality ((Lichtenstein et al. 1973; Choung et al. 2013), but see (Finlayson et al. 1975)). Our study offers a glimpse at the potentially antagonistic effect of glyphosate on spinosad efficacy: plants treated with glyphosate exhibited higher levels of herbivory damage in the presence of spinosad. This result is in line with previous work showing that glyphosate application results in increased herbivory damage (Zhang and Baucom 2024), as well as with other studies that demonstrate an increase of herbivores on herbicide-treated plants (Dewar et al. 2000; Johnson and Baucom 2022). Given that glyphosate can alter herbivory damage patterns (*e.g.*, hole-feeding vs. margin-feeding) in this system (Zhang and Baucom 2024), further work is necessary to determine whether spinosad may affect the community composition of insect herbivores.

In conclusion, our study examined plant resistance in different environmental contexts as mediated by glyphosate and spinosad, showing that spinosad modified the expression of glyphosate resistance by increasing it, thereby indirectly improving the fitness of exposed plants. Additionally, spinosad may alter the strength of selection acting on glyphosate resistance, potentially influence the evolutionary trajectory of glyphosate resistance. As plants in nature rarely face challenging conditions in isolation, it is important to examine the effects of complex environments on resistance traits to predict how plants may adapt in real time. Using factorial experiments to evaluate fitness effects across multiple environmental contexts and stressors offers a powerful way to understand how plants respond to shifting, multifaceted environments.

