## Supplemental Materials for "Insecticide increases glyphosate resistance in *Ipomoea purpurea*"

**SUPPLEMENTAL INFORMATION**

**Insecticide may alter the evolution of glyphosate resistance in *Ipomoea purpurea***

**Supplemental Figures**

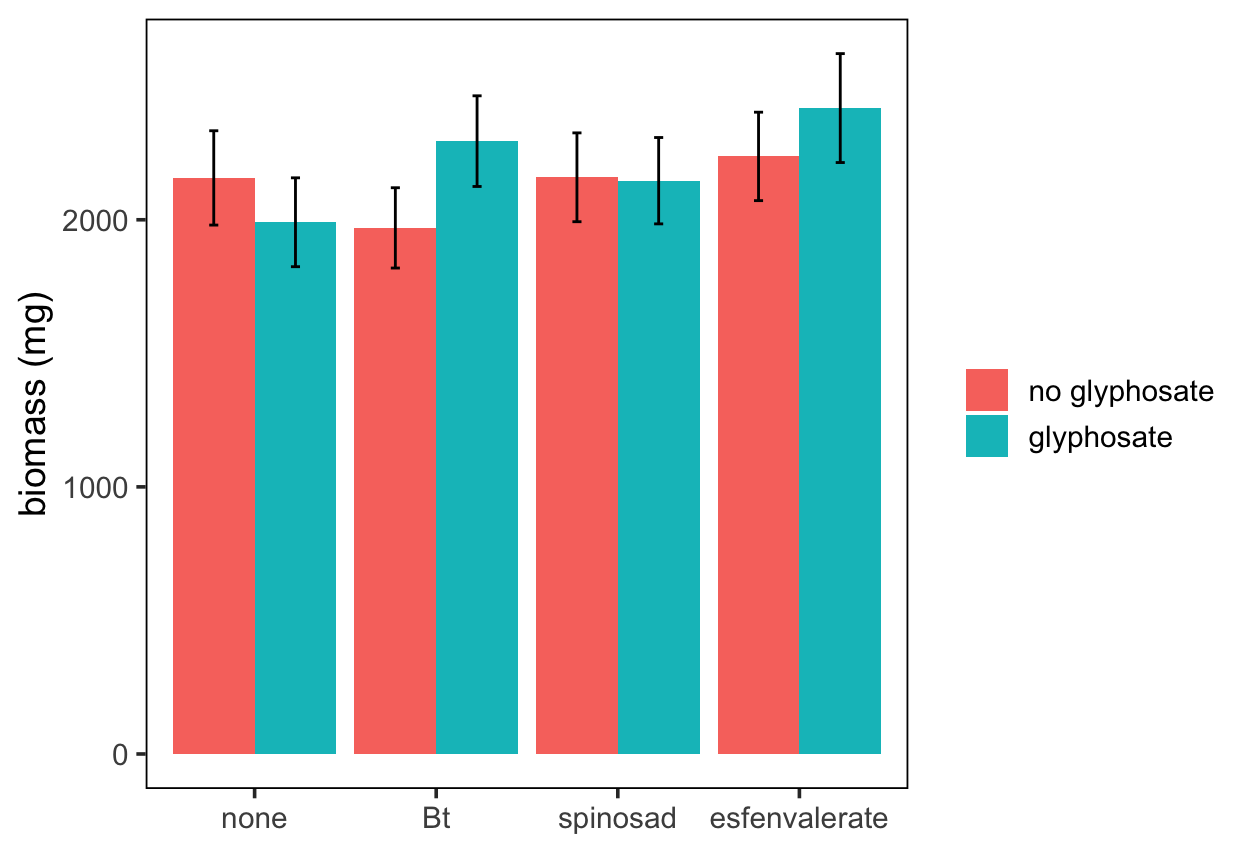

**Figure S1.** Biomass (mg) of *Ipomoea purpurea* plants treated with four insecticide treatments—none, Bt, spinosad, and esfenvalerate—in the presence or absence of glyphosate treatment.

Prior to the field study, in a separate experiment, we conducted a growth room study to determine which insecticide would not affect the performance of *I. purpurea* and would not interact with glyphosate, testing the effects of the insecticides Bt (*Bacillus thuringiensis* subsp. *kurstaki*), spinosad, and esfenvalerate. 40 *I. purpurea* seeds from 10 randomly-selected maternal lines from the US Midwest and South (Kuester et al., 2015) were scarified and planted in cone-tainers in trays, evenly divided in each of the following treatments: control, glyphosate, Bt, Bt + glyphosate, spinosad, spinosad + glyphosate, esfenvalerate, esfenvalerate + glyphosate—for a total of 50 seeds in each treatment, and 400 seeds overall. The growth rooms were on 12-hour light cycles and kept at 24-27°C. The number of leaves was measured six weeks after germination, and insecticides started being applied weekly at the recommended field dose for a total of four rounds. Glyphosate was applied at 0.84 kg ai/ha seven weeks after germination. Eleven weeks after germination, plants were clipped at soil level, collected in individual paper bags, dried for 48 hours at 60°C, then weighed for aboveground biomass.

To determine if the insecticides and glyphosate had an effect on biomass, we used a log transformation on the response variable, biomass. We then constructed the following linear model, wherein we tested for an interaction between insecticide and glyphosate, leaf count was included as a fixed effect covariate, and maternal line was a random effect:

biomass ~ (insecticide ✕ glyphosate) + leaf count + 1|maternal line

We found no evidence of a significant interaction between insecticide and glyphosate on biomass (F_1, 372_ = 2.12, p = 0.097), nor was there an effect of glyphosate alone (F_1, 372_ = 0.45, p = 0.502; Table S10). The type of insecticide, however, had an effect (F_1, 372_ = 8.25, p < 0.001). While Bt (t_372.55_ = 0.07, p = 0.942) and spinosad (t_372.42_ = 1.47, p = 0.143) did not impact biomass, esfenvalerate (t_372.60_ = 2.70, p= 0.007) had a positive effect on biomass. Additionally, there was an interaction between Bt and glyphosate that increased biomass (t_372.28_ = 2.51, p= 0.012). Given these results, we chose to use spinosad in the field experiment because unlike Bt and esfenvalerate, spinosad did not directly affect biomass, nor did it interact synergistically with glyphosate.

**Supplemental Tables**

**Table S1.** Linear mixed model for the effects of glyphosate and spinosad treatments on herbivory damage, with leaf count (as a proxy for plant size) and rust fungus included as fixed covariates. Family line and block were included as random effects and evaluated using the χ^2^ test of difference. Bolded p-values denote statistical significance, and italicized p-values show marginal significance.

| herbivory damage ~ spinosad + glyphosate + leaf count + rust + 1\|family + 1\|block | | | |
| --- | --- | --- | --- |
| **fixed effect** | **df** | **F** | **p** |
| spinosad | 1 | 198.94 | **<0.001** |
| glyphosate | 1 | 4.90 | 0.027 |
| leaf count | 1 | 10.89 | **0.001** |
| rust | 1 | 3.40 | *0.065* |
| **random effect** | **df** | **𝟀2** | **p** |
| family | 1 | 0.25 | 0.614 |
| block | 1 | 0.75 | 0.385 |

**Table S2.** Linear mixed model assessing the effect of glyphosate on herbivory damage in the presence and absence of spinosad, with leaf count (as a proxy for plant size) and rust fungus included as fixed covariates. Family line and block were included as random effects and evaluated using the χ^2^ test of difference. Bolded p-values denote statistical significance and italicized p-values denote marginal significance.

| herbivory damage ~ glyphosate + leaf count + rust + 1\|family + 1\|block | | | |
| --- | --- | --- | --- |
| **fixed effect** | **df** | **F** | **p** |
| *In the presence of spinosad* | | | |
| glyphosate | 1 | 5.32 | **0.021** |
| leaf count | 1 | 3.97 | **0.047** |
| rust | 1 | 1.79 | 0.182 |
| *In the absence of spinosad* | | | |
| glyphosate | 1 | 1.18 | 0.278 |
| leaf count | 1 | 8.64 | **0.003** |
| rust | 1 | 1.83 | 0.176 |
| **random effect** | **df** | **𝟀2** | **p** |
| *In the presence of spinosad* | | | |
| family | 1 | 3.29 | 0.070 |
| block | 1 | 0 | 1.000 |
| *In the absence of spinosad* | | | |
| family | 1 | 0 | 1 |
| block | 1 | 3.57 | *0.059* |

**Table S3.** Linear mixed model assessing the effect of spinosad treatments on glyphosate resistance, with leaf count (as a proxy for plant size) and rust fungus included as fixed covariates. The interaction between spinosad treatment and family line as well as block were included as random effects and evaluated using the χ^2^ test of difference. Bolded p-values denote statistical significance.

| glyphosate resistance ~ spinosad + leaf count + rust + (1 + spinosad\|family) + 1\|block | | | |
| --- | --- | --- | --- |
| **fixed effect** | **df** | **F** | **p** |
| spinosad | 1 | 82.17 | **<0.001** |
| leaf count | 1 | 84.99 | **<0.001** |
| rust | 1 | 12.32 | **<0.001** |
| **random effect** | **df** | **𝟀2** | **p** |
| 1 + spinosad\|family | 2 | 1.23 | 0.540 |
| block | 1 | 30.88 | **<0.001** |

**Table S4.** Linear mixed model assessing genetic variation for glyphosate resistance in the overall study and within spinosad-present and -absent environments. Family line, leaf count (as a proxy for plant size), and rust fungus were included as fixed effects, while block was included as a random effect. Bolded p-values denote statistical significance and italicized p-values show marginal significance.

| glyphosate resistance ~ family + leaf count + rust + 1\|block | | | |
| --- | --- | --- | --- |
| **fixed effect** | **df** | **F** | **p** |
| *Overall* | | | |
| family | 44 | 1.72 | **0.003** |
| leaf count | 1 | 66.56 | **<0.001** |
| rust | 1 | 7.72 | **0.006** |
| *In the presence of spinosad* | | | |
| family | 44 | 1.73 | **0.005** |
| leaf count | 1 | 11.39 | **0.001** |
| rust | 1 | 9.79 | **0.002** |
| *In the absence of spinosad* | | | |
| family | 44 | 1.77 | **0.004** |
| leaf count | 1 | 2.88 | *0.091* |
| rust | 1 | 2.41 | 0.122 |
| **random effect** | **df** | **𝟀2** | **p** |
| *Overall* | | | |
| block | 1 | 21.19 | **<0.001** |
| *In the presence of spinosad* | | | |
| block | 1 | 0 | 1.000 |
| *In the absence of spinosad* | | | |
| block | 1 | 123.38 | **<0.001** |

**Table S5.** Variance of glyphosate resistance attributable to family line. The effect of spinosad on variance partitioning was tested using the following model: glyphosate resistance ~ spinosad + leaf count + rust + (1 + spinosad|family) + 1|block. Variance components include among-family variance within each spinosad environment, among-block variance, and residual variance. Intraclass correlation coefficients (ICCs) represent the proportion of total variance attributable to family line within each spinosad environment. Point estimates and 95% confidence intervals were calculated using parametric bootstrapping (n = 1000 simulations).

| **parameter** | **estimate** | **95% CI lower** | **95% CI upper** |
| --- | --- | --- | --- |
| intercept | 0.084 | 0.021 | 0.169 |
| slope | 0.009 | 0.000 | 0.112 |
| spinosad-present | 0.084 | 0.021 | 0.169 |
| spinosad-absent | 0.038 | 0.005 | 0.102 |
| among-block variance | 0.093 | 0.000 | 0.488 |
| residual variance | 0.656 | 0.574 | 0.728 |
| spinosad-present ICC | 0.101 | 0.028 | 0.200 |
| spinosad-absent ICC | 0.049 | 0.007 | 0.130 |

**Table S6.** Linear mixed model assessing genetic variation for herbivory resistance in the overall study and within glyphosate-present and -absent environments. Family line, leaf count (as a proxy for plant size), and rust fungus were included as fixed effects, while block was included as a random effect. Bolded p-values denote statistical significance.

| herbivory resistance ~ family + leaf count + rust + 1\|block | | | |
| --- | --- | --- | --- |
| **fixed effect** | **df** | **F** | **p** |
| *Overall* | | | |
| family | 44 | 0.59 | 0.984 |
| leaf count | 1 | 12.22 | **0.001** |
| rust | 1 | 0.58 | 0.448 |
| *In the presence of glyphosate* | | | |
| family | 44 | 0.57 | 0.987 |
| leaf count | 1 | 0.92 | 0.339 |
| rust | 1 | 1.42 | 0.234 |
| *In the absence of glyphosate* | | | |
| family | 44 | 1.04 | 0.414 |
| leaf count | 1 | 7.56 | **0.006** |
| rust | 1 | 0.36 | 0.548 |
| **random effect** | **df** | **𝟀2** | **p** |
| *Overall* | | | |
| block | 1 | 4.86 | **0.027** |
| *In the presence of glyphosate* | | | |
| block | 1 | 0 | 1.000 |
| *In the absence of glyphosate* | | | |
| block | 1 | 8.74 | **0.003** |

**Table S7.** Variance of herbivory resistance attributable to family line. The effect of glyphosate on variance partitioning was tested using the following model: herbivory resistance ~ glyphosate + leaf count + rust + (1 + glyphosate|family) + 1|block. Variance components include among-family variance within each glyphosate environment, among-block variance, and residual variance. ICCs represent the proportion of total variance attributable to family line within each glyphosate environment. Point estimates and 95% confidence intervals were calculated using parametric bootstrapping (n = 1000 simulations).

| **parameter** | **estimate** | **95% CI lower** | **95% CI upper** |
| --- | --- | --- | --- |
| intercept | 0.001 | 0.000 | 0.078 |
| slope | 0.010 | 0.000 | 0.163 |
| glyphosate-present | 0.001 | 0.000 | 0.078 |
| glyphosate-absent | 0.004 | 0.000 | 0.084 |
| among-block variance | 0.030 | 0.000 | 0.159 |
| residual variance | 0.935 | 0.811 | 1.032 |
| glyphosate-present ICC | 0.001 | 0.000 | 0.078 |
| glyphosate-absent ICC | 0.004 | 0.000 | 0.083 |

**Table S8.** ANCOVA assessing the effect of spinosad on the selection gradients for glyphosate resistance at the family level in the two glyphosate-present environments. Relative fitness and standardized glyphosate resistance were calculated within-environment. Bolded p-values denote statistical significance and italicized p-values show marginal significance.

| relative fitness ~ glyphosate resistance + spinosad + glyphosate resistance * spinosad | | | |
| --- | --- | --- | --- |
| **fixed effect** | **df** | **F** | **p** |
| glyphosate resistance | 1 | 29.27 | **<0.001** |
| spinosad | 1 | 0.00 | 1.000 |
| glyphosate resistance * spinosad | 1 | 2.92 | *0.091* |

**Table S9.** Nonlinear selection analyses of glyphosate and herbivory resistance in each of the four treatment environments. Correlational selection models took the following format: relative fitness ~ glyphosate resistance × herbivory resistance + glyphosate resistance + herbivory resistance + glyphosate resistance^2^ + herbivory resistance^2^. Quadratic (i.e., stabilizing/disruptive) selection models for each individual form of resistance regressed relative fitness on both the linear and quadratic terms. Here, we only report the results of the nonlinear regression terms. Selection gradients and associated standard errors have been doubled from the values obtained from the linear regression. Italicized p-values show marginal statistical significance.

| **Trait** | **γ** | **SE** | **F** | **df** | **p** |
| --- | --- | --- | --- | --- | --- |
| *glyphosate-present, spinosad-absent* | | | | | |
| glyphosate resistance | 0.08 | 0.08 | 0.99 | 1 | 0.325 |
| herbivory resistance | -0.03 | 0.07 | 0.21 | 1 | 0.647 |
| *glyphosate-absent, spinosad-absent* | | | | | |
| glyphosate resistance | 0.003 | 0.03 | 0.02 | 1 | 0.902 |
| herbivory resistance | 0.01 | 0.02 | 0.15 | 1 | 0.698 |
| *glyphosate-present, spinosad-present* | | | | | |
| glyphosate resistance | -0.10 | 0.1 | 1.13 | 1 | 0.295 |
| herbivory resistance | 0.02 | 0.07 | 0.06 | 1 | 0.807 |
| *glyphosate-absent, spinosad-present* | | | | | |
| glyphosate resistance | -0.04 | 0.03 | 1.60 | 1 | 0.210 |
| herbivory resistance | -0.03 | 0.02 | 1.48 | 1 | 0.230 |
| **Environment** | **correlational selection gradient** | **SE** | **F** | **df** | **p** |
| glyphosate-present, spinosad-absent | 0.21 | 0.08 | 6.45 | 1 | **0.015** |
| glyphosate-absent spinosad-absent | 0.02 | 0.03 | 0.26 | 1 | 0.611 |
| glyphosate-present, spinosad-present | -0.18 | 0.13 | 1.79 | 1 | 0.189 |
| glyphosate-present, spinosad-present | -0.02 | 0.03 | 0.25 | 1 | 0.622 |

**Table S10.** Linear mixed model testing the effects of glyphosate and three insecticides (Bt, spinosad, and esfenvalerate) on biomass in the preliminary insecticide selection growth room experiment. Leaf count was included as a fixed-effect covariate, while maternal line was included as a random effect and evaluated using the χ^2^ test of difference. Bolded p-values denote statistical significance, while italicized p-values show marginal statistical significance.

| biomass ~ insecticide * glyphosate + leaf count + (1\|maternal line) | | | |
| --- | --- | --- | --- |
| **fixed effect** | **df** | **F** | **p** |
| insecticide | 3 | 8.25 | **<0.001** |
| glyphosate | 1 | 0.45 | 0.502 |
| leaf count | 1 | 1107.30 | **<0.001** |
| insecticide * glyphosate | 3 | 2.12 | *0.097* |
| **random effect** | **df** | **𝟀2** | **p** |
| maternal line | 1 | 2.46 | 0.117 |
